# A transcriptional map of human tonsil architecture: beyond the sum of (single cell) parts

**DOI:** 10.1101/2025.09.08.674825

**Authors:** Helena L. Crowell, Laura Llaó-Cid, Gerard Frigola, Samuel Gunz, Irene Ruano, Patricia Lorden, Max Ruiz, Marta Kulis, José Ignacio Martin-Subero, Holger Heyn, Elias Campo, Anna Pascual-Reguant

**Affiliations:** Centro Nacional de Análisis Genómico (CNAG), Barcelona, Spain; Institut d’Investigacions Biomèdiques August Pi i Sunyer (IDIBAPS), Barcelona, Spain; Hematopathology Section, Pathology Department, Hospital Clínic, Barcelona, Spain; Institute of Molecular Life Sciences, University of Zurich, Zurich, Switzerland; Swiss Institute of Bioinformatics (SIB), Zurich, Switzerland; Centro de Investigación Biomédica en Red de Cáncer (CIBERONC), Madrid, Spain; Universitat de Barcelona (UB), Barcelona, Spain; Institución Catalana de Investigación y Estudios Avanzados (ICREA), Barcelona, Spain

## Abstract

The tonsil is a highly compartmentalized organ in which different microanatomical structures orchestrate designated (immune) functions. We use this already well-studied tissue to survey spatial molecular imaging data (CosMx SMI) for studying immune responses in native tissue context; and, to demonstrate the advantages of SMI for faithfully recapitulating cellular composition in direct comparison with single-cell RNA sequencing. While SMI data still poses many analytical challenges and lacks standardization, we established a versatile analysis pipeline focused on the profitable particularities of these data: considering organization (microenvironment), interactions (signaling), and function (higher-order structures) across scales.

Specifically, we resolve ~ 2M cells into 52 subpopulations across immune and, in particular, structural compartments. Various spatial niches partition tonsillar tissue into architecturally and functionally distinct regions, which we characterize through cell-cell colocalization and communication analyses, while performing various non-standard analyses at the level of spatial features. These topological readouts may help elucidate where certain immunological processes occur (e.g., class switch recombination); and, where signaling pathways are active (e.g., TNF and galectin, which have been implicated in diverse lymphomas).

In all, we provide an analytical framework for Spatial Immunology, and showcase alternative views that such techniques and concomitant computational approaches can bring on tissue composition and architecture.

## Introduction

Multi-cellular systems orchestrate function through an interplay of their molecular components and structural organization (e.g., tissue composition and architecture). For the immune system, cellular positioning is critical for both cell homeostasis and generation of protective responses during infection and vaccination^[1,2]^. The position of the palatine tonsils at the gateway of both the respiratory tract and the digestive tract leads to constant exposure to microorganisms including commensal bacteria and pathogenic viruses. As a secondary lymphoid organ (SLO), the tonsil is a highly compartmentalized and spatially structured tissue, in which every microanatomical region fulfills a specific (immune) function. Linked to that, each region is enriched in distinct structural (epithelia, fibroblasts, and endothelia) and immune cell types. The tonsil compartments are thus named: B cell follicles, the T cell zone, connective tissue septum, and the epithelium, subdivided into surface and reticular crypts.

Our previous work has identified rare immune cell subtypes (ILCs) and their niches in the tonsil using a highly multiplexed imaging technique coupled with a customized analysis pipeline^[3]^. More recently, we generated an atlas of cells in the human tonsil^[4]^ using various modalities, including single-cell RNA sequencing (scRNA-seq) and sequencing-based spatial transcriptomics, which was limited by its lateral resolution and unable to distinguish single cells in this regard. Nowadays, imaging-based spatial transcriptomics technologies can profile tissues at molecular resolution while retaining physical coordinates of both targets and cells; as of recent, they are able to resolve the protein-coding transcriptome^[5,6]^. However, analyzing data from such assays remains challenging, and lacks standardization compared to scRNA-seq^[7,8]^. It is, for instance, not clear whether the statistical assumptions that underlie tools developed for scRNA-seq data are met. Yet, these data have the potential to expand our biological understanding by viewing tissues as more than the sum of their single cell parts.

Building upon the Tonsil Cell Atlas^[4]^, we set out to demonstrate how CosMx Spatial Molecular Imaging (SMI) data can yield fresh biological insights into this already well-known and -characterized tissue^[9,10,3,4]^. We compare the absolute and relative abundance of cells profiled for each lineage in scRNA-seq and CosMx SMI data, and explore higher levels of hierarchical functional organization by stratifying sections into distinct spatial niches that represent well-defined histological tonsil regions. At the niche- and subpopulation-level, within and across sections, we interrogate cell-cell communication modeling 181 signaling interactions. Through unit-based analyses, we quantify expression gradients within germinal centers (GCs), their composition across sections, as well as how cells localize within, and differ across apical-basal/crypt-surface type epithelium. Treating GCs as spatially segregated units also let us study class switch recombination on a per-GC basis, and investigate potential correlates (e.g., presence of contextually relevant subpopulations and cytokines). In all, we believe this study highlights the power of SMI-based assays and spatial-centric analyses to unravel the real composition and cellular organization of tissues, in particular, of such a crucial immunological organ as the tonsil.

## Results

### Imaging-based ST captures tissue complexity more faithfully than scRNA-seq

Here, we acquired imaging-based spatial transcriptomics data (CosMx SMI^®^, Bruker) on three palatine tonsil sections (C1, A1, and A2) using the Universal Cell Characterization Panel, which includes 980 RNA targets, 10 negative probes (to quantify non-specific ISH probe hybridization), and 197 blank codes (to quantify misidentification of reporter readout), as well as immunofluorescence (IF) markers for B2M/CD298 (membrane), DAPI (nuclei), CD45 (immune), CD68 (macrophages) and PanCK (epithelia); see Fig. 1a and Fig. S1. Section C1 came from a child, sections A1 and A2 from different adults. We called 2,089,984 cells across 659 fields of view (FOVs), amounting to a total scanning area of ~ 1,73cm^2^ (Table S1).

**Figure 1:**
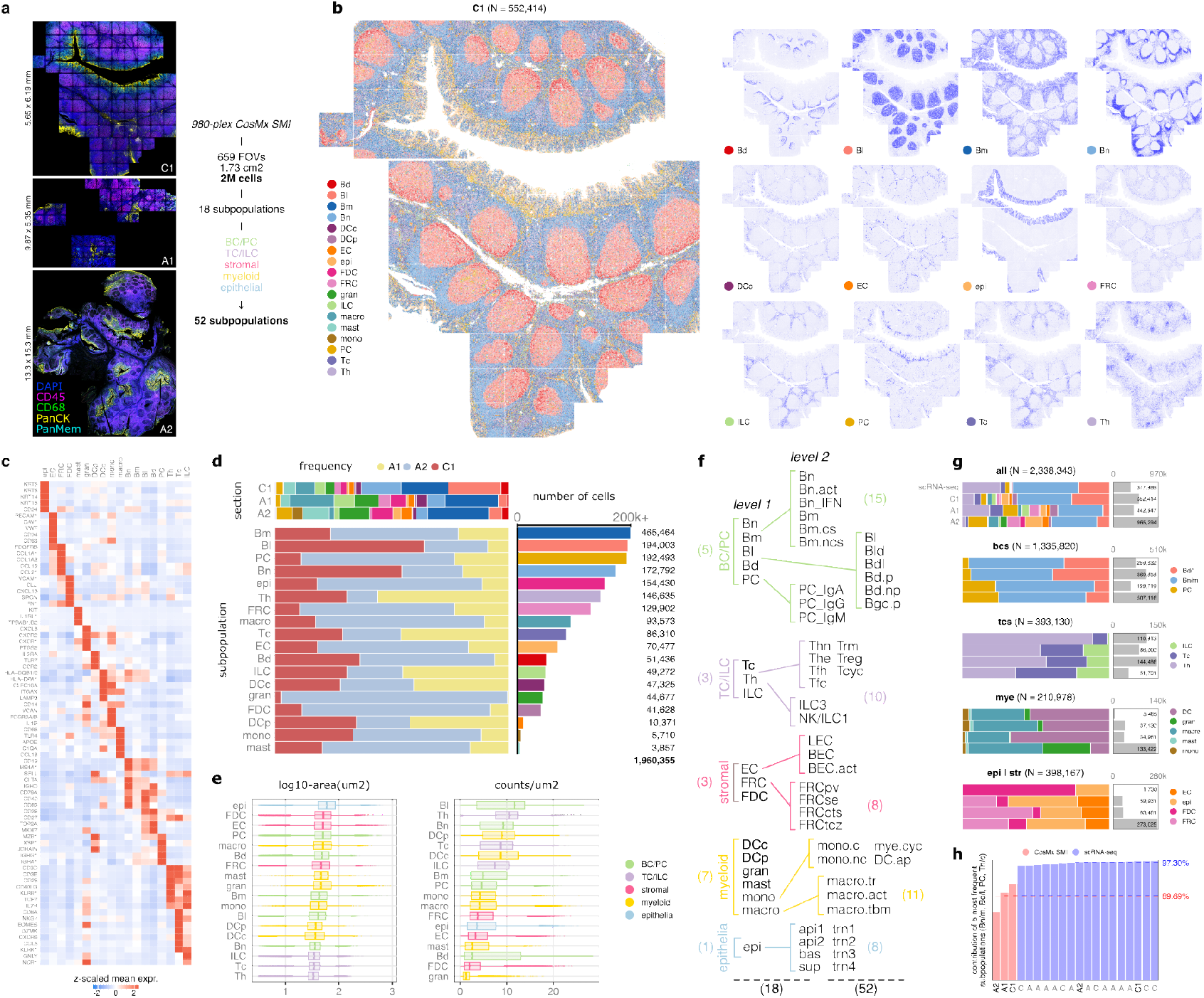
Transcriptional and compositional tissue characterization. **(a)** Schematic data summary including immunofluorescence composite images, the total number of cells and fields of view (FOVs) acquired, their cumulative scanning area, as well as the number of subpopulations annotated. **(b)** Spatial plot for section C1 with cells colored by subpopulation; and, spatial plots highlighting selected subpopulations individually (included are all subpopulations with *>*5k cells except for macrophages that were evenly distributed). **(c)** Heatmap displaying the mean expression of selected marker genes, z-scaled across subpopulations (data across all sections). **(d)** Subpopulation frequencies across sections and vice versa, as well as total number of cells by subpopulation. **(e)** Comparison of cellular areas and target counts per area across subpopulations; y-axes ordered by median. **(f)** Schematic of hierarchical clustering employed to arrive at high-resolution annotations. Subpopulations from each compartment were pooled and subclustered using distinct feature selections, yielding a total of 18 and 52 subpopulations at level 1 and 2, respectively; subpopulation labels that are in bold at level 1 are also present at level 2. **(g)** Comparison of absolute and relative subpopulation abundances between data acquired through scRNA-seq and CosMx SMI. **(h)** Relative contribution of subpopulations that are most frequent in Massoni-Badosa *et al*.^[4]^ across scRNA-seq and CosMx SMI samples; C/A = child/adult, dashed lines indicate averages, labels of paired samples are highlighted in black.

Quantifying the number of RNA counts per cell as a function of their physical distance to each FOV border, we observed a sharp decline in counts near borders, followed by onset of a plateau around the tissue-wide average (Fig. S2a). To mitigate potential artifacts in downstream analyses, we thus filtered out cells too close to any FOV border, as well as cells with few features and counts (in total and per area); cf. Fig. S2b. Notably, we observed distinct counts as a function of distance to top/left and bottom/right FOV borders, respectively. This observation was in line with the corner effect observed from IF stains, with reduced signal in top-left FOV corners (cf. Fig. S1). Both border and corner effects were stronger in the first dataset produced (sections C1 and A1; see Methods). Overall, we retained 1,960,355 cells (~ 93,8%) for downstream analysis (Table S1).

Semi-supervised clustering with reference profiles derived from Massoni-Badosa *et al*.^[4]^ yielded 18 subpopulations: plasma cells -PC-, naive -Bn- and memory -Bm-B cells, germinal center B cells (GCBCs) of the light -Bl- and dark zone -Bd-, helper -Th- and cytotoxic T cells -Tc-, innate lymphoid cells -ILC-, endothelial cells -EC-, follicular dendritic cells -FDC-, fibroblast reticular cells -FRC-, macrophages -macro-, monocytes -mono-, mast cells, granulocytes -gran-, conventional -DCc- and plasmacytoid -DCp-dendritic cells, as well as epi(thelia). These subpopulations exhibited a marked spatial organization (Fig. 1b; see also Fig. S3, Fig. S4, and Fig. S5), and had distinct transcriptional signatures that included marker genes established in the scientific literature (Fig. 1c). Subpopulations comprised ~ 4-465k cells, with mast cells being the rarest, and Bm the most abundant (Fig. 1d). All were represented across sections, albeit in varying proportions; most notably, Bn and Bl subpopulations that make up B cell follicles (mantle and GC light zone, respectively) were much more frequent in section C1, in line with an age-related decline of the GC reaction that is characterized by a smaller light zone, and accompanied by a change in the follicle’s spatial organization^[11]^.

We next quantified the area and target counts per area across different subpopulations (Fig. 1e). Epithelia were largest, followed by stromal subtypes and PCs, then B and myeloid cells; T lymphocytes and ILCs were the smallest. We detected a median of ~ 1-12 counts per um2, with gran, FDCs, and Bd being at the lower end, followed by other myeloid, stromal, epithelial subpopulations, as well as Bm and PCs; DCs, T cells and ILCs, as well as Bn and Bl were at the upper end.

Pooling epithelial -epi-, myeloid -mye-, stromal -str-, plasma and B cell -bcs-, as well as T cell and ILC subpopulations -tcs-, we subclustered them separately to obtain 52 high-resolution subpopulations (Fig. 1f, Fig. S6, Table S2; see Methods). In the BC/PC compartment (number of subpopulations N=15), we resolved three PC subtypes that differed in their Ig signature, six GC, as well as three Bn and Bm subtypes each. In the TC/ILC compartment (N=10), we identified various helper, cytotoxic, effector, tissue-resident memory and follicular T cell subtypes, as well as NK/ILC1 and ILC3. The myeloid compartment (N=11) consisted of non-/classical monocytes, three DC and macro subtypes each, gran and mast cells, as well as a cycling subpopulation.

The stromal compartment (N=8) comprised FDCs located in the follicle, four FRC subpopulations distributed differently across regions, two blood and one lymphatic EC subtype(s). Tonsillar FRCs were named according to their location (namely, T cell zone -tcz-, connective tissue septum -cts-, subepithelium -se-, and perivascular -pv-; cf., Fig. S7) that was accompanied by distinct transcriptional profiles associated therewith, as previously shown in other SLOs^[12,13,14,15]^. Within the epithelial compartment (N=8), we delineated two apical, basal, suprabasal, and four transitional subtypes. Despite limited plexity, key canonical marker genes were exclusive to each of these high-resolution subpopulations (Fig. S6).

Next, we compared relative subpopulation abundances to scRNA-seq reference atlas data^[4]^ from3 adults and 7 children, including matched samples for sections C1 and A1 (Fig. 1f,g). The five most frequent subpopulations in the scRNA-seq data were Bn/m, Bd/l, PC, Th and Tc; together, these accounted for ~ 97% of cells per scRNA-seq sample (Fig. 1h). By contrast, they amounted to only ~ 70% per section acquired through CosMx SMI. The number of cells in our data were up to two orders of magnitude higher per compartment, although this will arguably depend on the number of FOVs. Here, an average ~ 3k cells were segmented per FOV (cf. Table S1). As such, 2-4 FOVs would be sufficient to surpass the 5-10k cells per sample acquired in a standard scRNA-seq experiment (by comparison, 200 FOVs per slide are usually used for CosMx experiments with the 1K and 6K panel parallelizing 4 and 2 slides, respectively). There was a complete loss of ECs and FRCs in the scRNA-seq data, which accounted for ~ 50% of the structural compartment in our sections. ILC were of similar abundance as Tc in our data, but very rare across scRNA-seq samples.

In general, scRNA-seq mainly captured follicular subpopulations – over-representing GCBCs, FDCs, and Th cells – and, in turn, reflects less heterogeneity across other compartments, in particular, of tissue-resident and stromal cells.

### Imaging-based ST resolves architecture and functional designation of distinct tissue regions

The communication between immune cells within particular SLO compartments is organized by fibroblasts that are commonly referred to as fibroblast reticular cells (FRCs)^[16,17]^. Specialized FRC subsets form the scaffold and create the milieu for lymphocyte guidance, activation and differentiation in distinct specialized niches^[18,14]^. We aimed at mapping these niches at the cellular and molecular level.

Quantifying local neighborhoods of subpopulations across sections unveiled distinct colocalization patterns (Fig. 2a); e.g., a follicular module (GCBCs, Tfh, FDC, macro.tbm/act), an epithelial module (gran, mono.c, epi.x), a T cell module (DCc alongside different T cells and FRCtcz), and a highly vascularized module, rich in myeloid cells. Interestingly, tissue-resident memory T cells -Trm-, Bm.cs, ILC and DC subpopulations, were found close to both, the T cell module and a B cell module, composed of FRCse, Bn and Bm. Based on these neighborhoods, we clustered cells into similarly composed microenvironments, thus compartmentalizing the tissue in a data-driven manner as opposed to (manual) histopathology-based annotation (see Methods).

**Figure 2:**
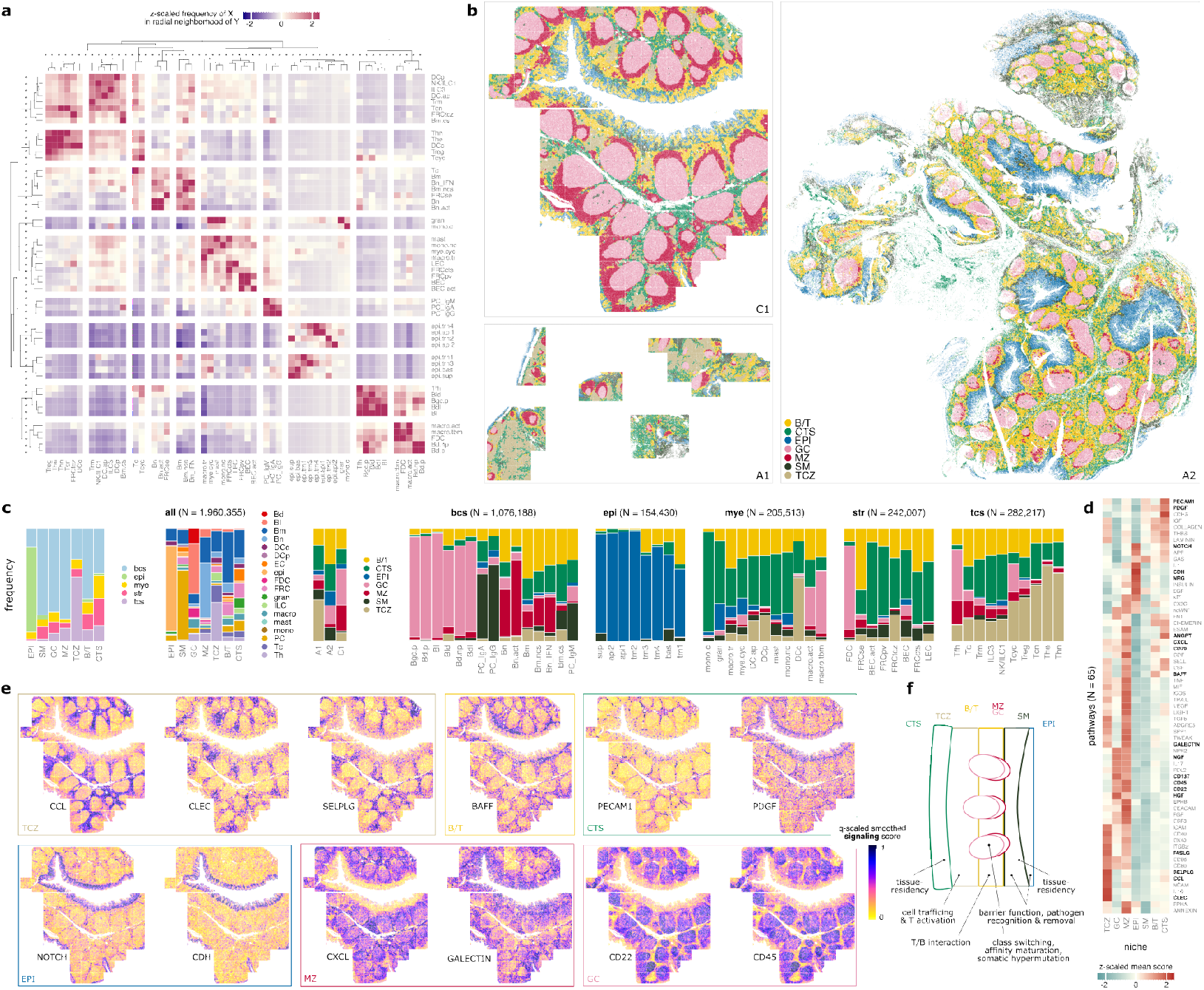
Niche analysis. **(a)** Heatmap quantifying cell-cell colocalization between subpopulations (data across all sections); rows and columns ordered by hierarchical clustering, and split by (*k*=10)-means clustering. Bin values correspond to the proportion of cells from subpopulations on the x-axis within the 20um radial neighborhood of subpopulations on the y-axis, averaged across cells, and z-scaled by row. **(b)** Spatial plots with cells colored by niche assignment: germinal center (GC) and mantle zone (MZ), B/T border, T cell zone (TCZ), connective tissue septum (CTS), epithelium (EPI) and submucosa (SM). **(c)** Frequency niches across sections (third panel) and of (fltr) compartments, low- and high-resolution subpopulations across niches; x-axes ordered by hierarchical clustering. **(d)** Heatmap of cell-cell communication pathways across niches; values correspond to sender and receiver signals averaged across all cells in a given niche, averaged with each other, and z-scaled across niches. **(e)** Spatial plots with cells colored by selected signaling pathways; sender and receiver values are each averaged across 20um radial neighborhoods, then with each other, and 0-1 scaled using lower/upper 1% quantiles as boundaries. **(f)** Schematic summarizing key functions of tonsillar niches. (^*^Besides the epithelium (EPI) and follicles (GC/MZ), niches are mixed and not in fact layered).

Considering frequency profiles and spatial organization, we annotated seven niches that aligned with known tissue architecture, namely: germinal center (GC), mantle zone (MZ), T cell zone (TCZ), B/T border, connective tissue septum (CTS), epithelium (EPI), and submucosa (SM) (Fig. 2b). Although all niches were represented across all sections, their relative abundances were highly variables due to both biological (age) and technical (FOV placement) differences (Fig. 2c). Around 50% of cells were follicular in section C1, of the TCZ in section A1, and of the B/T border and CTS in section A2, which was the most heterogeneous (and the largest tissue area analyzed).

The GC niche contained mostly Bd/l together with macro.act/tbm, FDCs, Tfh and Tcyc. The MZ was dominated by B cells (mainly Bn/.act, fewer Bm subtypes), alongside some T cells (all subtypes, mostly Tfh). Besides non-follicular B and all types of T cells, the TCZ included DCs (mostly conventional) as well as FRCtcz. Stromal and myeloid fractions representing different FRC and EC subpopulations were highest for the CTS, in accordance with its known high vascularization and extracellular matrix enrichment; in addition, the CTS housed a variety of T and ILC subsets alongside PCs, Bm and Bn IFN, mainly tissue-resident immune cells, as previously shown^[3]^. The B/T border mainly comprised B and T lymphocytes, but captured other subpopulations across all compartments in similar amounts, exceptions being epithelial and follicular subpopulations (Bd/l, macro.act/tbm, Tfh and FDCs). Notably, IgM+ PCs were equally distributed across B/T border, CTS, GC, and SM niches; by contrast, IgA+ and IgG+ PCs were much more frequent in the latter. The SM was particularly rich in PCs, found alongside Bm.cs as well as some stromal, less myeloid, and all but few T cells. Alongside epithelia, the EPI niche included some PCs and Bm, as well as a variety of myeloid subtypes excluding macro.act/tbm and DCc.

Lastly, we modeled cell-cell communication (CCC) at the single-cell level, considered 143 interactions across 38 pathways that were available through *CellChatDB*^[19]^ and for which all interaction partners were represented in our panel (using *COMMOT* ^[20]^; see Methods). Summarization at the niche-level highlighted specific pathways that dominate different microanatomical regions (Fig. 2d; see also Fig. S8); the spatial distribution of selected pathways is shown in Fig. 2e.

Follicular niches (GC and MZ) shared high HGF, NGF and CD137 (also known as TNFRSF9 or 4-1BB), all FDC-produced signals that promote GC formation and maintenance either by facilitating survival and function of FRCs^[21]^, GCBCs^[22,23]^, or Tfh cells. These niches also share CD45 and CD22, both controlling B cell receptor signaling in the follicles, and influencing persistence^[24]^ and output^[25]^ of the GC reaction. Particularly enriched in the MZ, GALECTIN signaling regulates B cell activation^[26,27]^ and survival; notably, it has also been implicated in the transformation and/or progression of various lymphomas^[28,29,30]^. CXCL pathway, although higher in the mantle, is ubiquitously expressed within GCs and interfollicular regions (TCZ and B/T border). The expression of different ligands by spatially restricted FRCs helps guide the localization of specific cell subsets into different tonsillar niches, thereby fostering particular cellular interactions and consequent functions^[31,32]^.

Within the CTS, we found increased angiopoietin (ANGPT) signaling, known to control vascular morphogenesis and homeostasis^[33]^. BAFF and CD70 signaling was higher in the T/B border, in line with their dual role on B and T cell lineages^[34,35]^, promoting the development and activation of follicular and MZ B cells, as well as Tfh. The TCZ showed an increase in CLEC, CCL, SELPLG and OX40 signaling. The first three play crucial roles in TCZ organization and function, promoting T cell and DC trafficking and interactions^[36,37,38]^. The latter promotes division and survival in both Th and Tc cells, augmenting the clonal expansion of effector and memory populations as they are being generated to antigen^[39]^. Complementarily, there was an upregulation of FASLG, a pathway involved in down-regulation of immune reactions (death/apoptotic signal), as well as in T cell-mediated cytotoxicity (co-stimulation)^[40,41]^. The EPI showed increased NOTCH, CDH, insulin, EGF, NRG, and IL1 signaling, all related to epithelial proliferation, barrier function maintenance, and regeneration in various tissues^[42,43,44,45]^; SM-related pathways included PDGF and GAS (see below for a more in-depth characterization of these niches).

Thus, imaging-based ST allowed us to map the precise cellular composition of, and to interrogate the signaling pathways that cells engage with in each tonsillar niche or compartment; ultimately, enabling a comprehensive study of how tissue architecture orchestrates tissue function (Fig. 2f).

### Unit-based analysis of germinal centers illuminates their composition, zonation, maturation, and efferocytic function

GCs are key immune structures that orchestrate humoral immunity. Their ultimate goal is the generation of i) plasma cells that can secrete high-affinity antibodies to fight infection; and, ii) memory cells that can mount rapid responses upon re-challenge^[46,47]^. To this end, GCs support distinct functions that occur in different regions: the dark zone (DZ) and the light zone (LZ)^[48]^. CXCL12+ reticular stromal cells guide CXCR4+ B cells to the DZ, where they proliferate and introduce mutations into their B cell receptor genes through somatic hypermutation (SHM). CXCL13+ FDCs guide CXCR5+ B cells to the LZ, where they undergo selection based on the strength of antigen binding and presentation to Tfh cells. CD40L-dependent Tfh co-stimulation prevents B cell death and induces Myc expression, driving re-entry into the cell cycle.

Based on the point pattern density of FDCs and GC B cells, we reconstructed GCs for sections C1 and A1 that were acquired together, identifying 71 GCs (using *sosta*; see Methods). For each GC, we also quantified how within-GC cells are positioned relative to nearby Bn cells that constitute the GC mantle (Fig. 3a,b, Fig. S9; see Methods). GCs were concordant with the known orientation of LZs towards the mucosal surface^[49]^, and contained ~ 200-12,000 cells (~ 3,127 on average). Their subpopulation composition was comparable within sections; however, compliant with an agerelated tonsillar involution^[50]^, GC B cells were less frequent in the adult sample (Fig. 3c).

**Figure 3:**
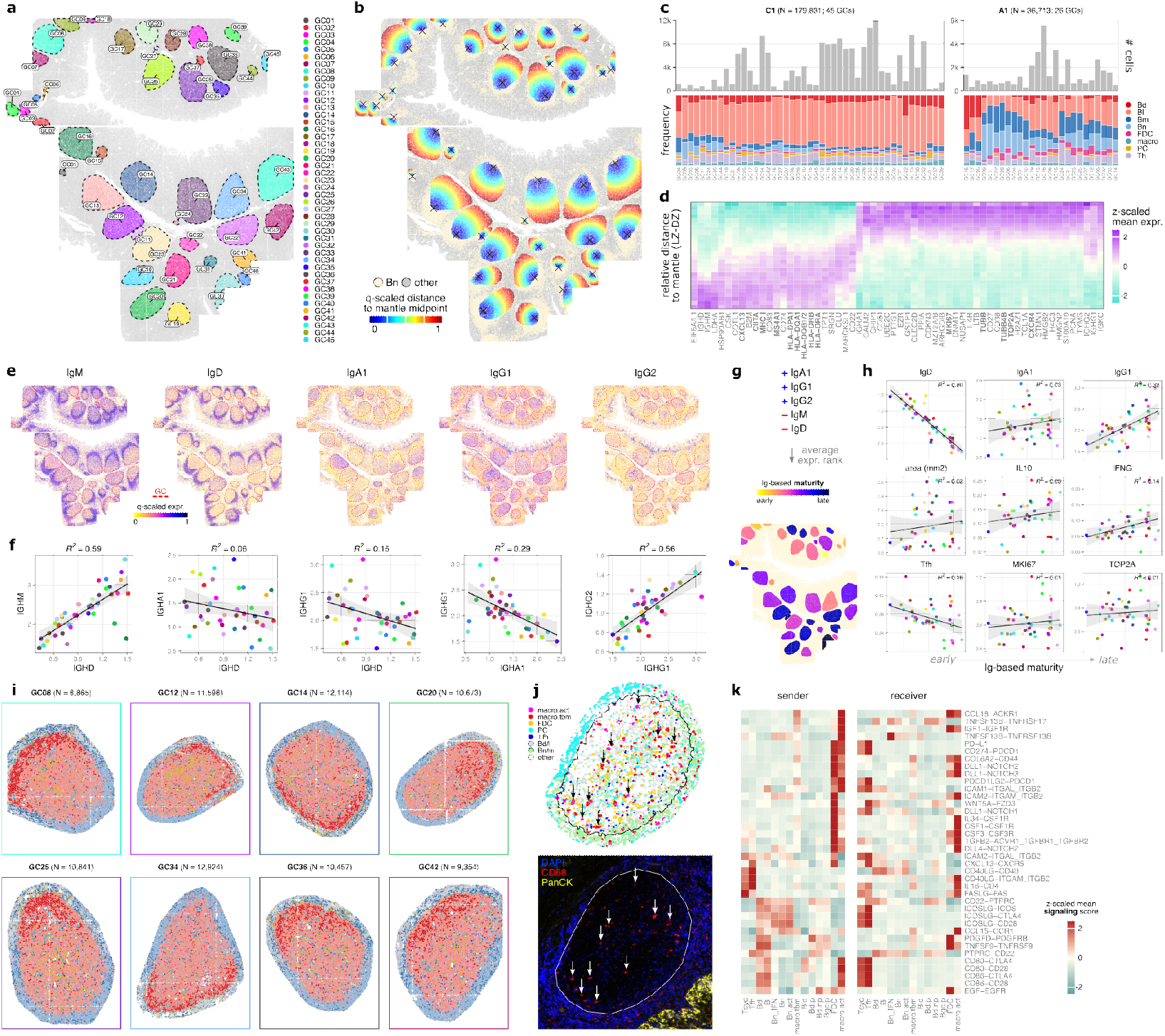
Germinal centers. **(a)** Spatial plot highlighting programmatically identified germinal centers (GCs); dashed outlines are concave hulls of selected cells (cf., Fig. S9). **(b)** Spatial plot with cells colored by their relative distance to the median coordinates (black crosses) of Bn (off white) that fall within a 50um polygonal expansion from a given GC; cells that are neither Bn nor in any GC are grayed out. **(c)** Absolute and relative abundance of key subpopulations across germinal centers (GCs); x-axes ordered by hierarchical clustering. **(d)** Expression heatmap binned by relative distance to respective reference anchors; included are the 40 genes with the highest value range across bins. **(e)** Spatial plots with cells color by expression of immunoglobulins (Ig’s); dashed red outlines are concave hulls of GCs. **(f)** Scatter plots comparing Ig levels across GCs; values correspond to mean expression across B cells (of any type) that fall within a given GC, but excluding the mantle. **(g)** Schematic illustrating Ig-based maturation scoring of GCs and corresponding spatial plot: Ig’s are ranked across GCs (by decreasing IgD/M, and increasing IgA1/G1/G2 expression), and the average rank across Ig’s is treated as GC maturity (low=early, high=late). **(h)** Scatter plots of maturity against Ig expression levels (averaged across B cells), surface area (in mm^2^), proportion of Tfh, as well as IL10/IFNG and MKI67/TOP2A expression (averaged across Tfh and Bd cells, respectively). Each point in (f,h) is a GC, color code according to (a). **(i)** Spatial plots for selected GCs colored by key subpopulations; included are cells within a 50um polygonal expansion of the structure’s concave hull (cf., Fig. S10). **(j)** Exemplary GC for section A2 highlighting key follicular subpopulations (left) alongside the corresponding IF image (right); black/white outlines are the GC selection boundary, arrows indicate selected CD68+ macro.act/tbm aggregates. **(k)** Heatmap of cell-cell communication interactions between GC-related subpopulations (section C1); sender and receiver scores were averaged by subpopulation, ranked by fold change between subpopulations, and the top-50 selected for visualization; axes are ordered by hierarchical clustering of ‘sender’ values.

In order to identify genes that – independent of cell types – exhibit a spatial association with the well-defined functional zonation of GCs, we quantified gene expression along the LZ-DZ axis, setting the mantle interface as an anchor point (see Methods). We indeed identified two distinct sets of genes that are relatively higher expressed in the dark and light zone, respectively (Fig. 3d). Genes associated with proliferation (MKI67, TUBB, TOP2A) and CXCR4 increase towards the DZ, while genes related to antigen presentation (HLAs, MHC I) and maturation (CD83, CIITA), together with CXCL13 increase towards the LZ, showing that tissue topology guides gene expression gradients beyond discrete cell types or regions.

After iterative rounds of SHM and selection, so-called affinity maturation, GC cells exit the GC, either as fully differentiated memory B cells, or as long-lived PCs that secrete immunoglobulins (Ig). In addition to SHM, B cells undergo Ig isotype switching, or class switch recombination (CSR), in response to antigen stimulation and co-stimulatory signals. CSR brings forth different Ig isotypes (from IgM and IgD production to either IgA, IgG, or IgE) without altering specificity for the immunizing antigen^[51]^. These two processes are distinct, having different mechanisms and outcomes, but are coordinated by the same enzyme within GCs: activation-induced cytidine deaminase (AID)^[52]^.

Tissue-wide expression of Ig isotypes (Fig. 3e) showed distinct spatial patterns that were in line with known regions for particular BC differentiation states. IgM and IgD were most highly expressed in the mantel zone that mainly consists of Bn; by contrast, IgG1/2 and IgA1 were restricted to GCs and the sub-epithelial layer, i.e., regions where PCs, activated and memory BCs are located. Within GCs (excluding their mantle), expression levels of IgM and IgD, as well as IgG1 and IgG2 were highly correlated (Fig. 3f). Although less pronounced, IgG1 and IgA1 were negatively correlated, indicating that the two Ig’s might be mutually exclusive between follicles. In connection herewith, we observed individual GCs that had comparatively high IgA1 but low IgG1/2 expression (e.g., GC22), and vice versa (e.g., GC35); cf., Fig. 3a,e. There was little association between IgD and IgG1, and none between IgD with IgA1.

Because CSR occurs in a fixed sequence, we hypothesized that differences in Ig expression at the GC level could be used as a proxy for follicle maturation. Specifically, we ranked the average expression of each Ig across GCs (by increasing value for IgA1/G1/G2, and decreasing value for IgD/M), and estimated each GC’s maturity as the average rank across Ig’s (Fig. 3g). Neither the expression of molecules (e.g., IL10, IFNG, TGFB1) nor the proportions of cell types (e.g., Tfh) known to drive class switching correlated with Ig-based maturation status (Fig. 3h). The expression of genes associated with proliferation (e.g., MKI67, TOP2A) in Bd subpopulations of GCs, as well as their surface area, also lacked association. Ranking based on a single Ig (e.g., only IgD or IgG1), or weighting ranks by position in the CSR sequence (i.e., {−1, −2, +3, +4, +5} for IgM/D/G1/A1/G2), did not significantly alter results. This lack of association (between Ig-based maturity and known CSR drivers) suggests that class switching occurs outside of GCs, supporting recent work by Roco *et al*.^[53]^ that challenged the classical view of CSR being a hallmark of GCs.

Beyond the B cell lineage, we identified a macrophage subpopulation (macro.tbm) largely found in GCs (Fig. 2c) that was prone to form cell aggregates (Fig. 3h) and strongly co-localized with FDCs, macro.act, and Bd (Fig. 2a). CD68 expression at the protein level validated the identity of these cells and their particular distribution pattern within GCs (Fig. 3j). These macrophages were characterized by the expression of classical macrophage markers, genes related to phagocytosis (e.g., SLC40A1, GPNMB^[54]^), as well as CLU and NUPR1, two key protective proteins known to inhibit various forms of cell death, thus promoting macrophage survival and anti-inflammatory functions^[55,56]^; Fig. S6.

Based on these results, we hypothesized this particular macrophage subset to represent tingible body macrophages (TBMs). These have previously been identified through imaging^[57,58]^ and shown to phagocytize apoptotic B cells that accumulate during the GC reaction, thus aiding to prevent secondary necrosis and autoimmune activation (in particular, Gurwicz *et al*.^[58]^ have shown that FDCs license TBMs to engulf apoptotic bodies). Even as part of a larger cohort, this macrophage subset could not be resolved in previous scRNA-seq analysis of these samples^[4]^, re-emphasizing the power of spatially-resolved data to detect rare cell populations with distinct spatial distributions.

CCC analysis of follicular subpopulations (Fig. 3k) confirmed key known interactions, such as costimulatory contact-dependent interactions of Tfh with Bl and Bn subsets through CD40LG-CD40 and ICOS-ICOSLG/CTLA4/CD28; between CD80/CD86 expressed by antigen presenting cells (light zone B cells and macro.tbm/act.) and CTLA-4/CD28, which play a crucial role in regulating Tfh development and function by either suppressing or promoting GC responses^[59,60,61]^; and, FDCs attracting Bl, Bn and Tfh through the chemokine CXCL13 and its cognate receptor CXCR5. Tfh are known to express high levels of PDCD1^[62]^, and its ligation with CD274 – constitutively expressed by follicular B cells – has been shown to control positioning and function of Tfh^[63]^. Instead of follicular B cells, we found FDCs as well as macro.tbm/act to be the main producers of CD274, and to communicate strongest with Tfh. We also detected GCBCs expressing TNFSF9 (signaling through TNFSRF9 on FDCs and macro.tbm/act), which has been implicated in lymphoma^[64,65]^ and, in mice, aberrant GC and autoantibody production^[66]^.

In summary, we have demonstrated how analyses that treat GCs as separate units (as opposed to a tissue-wide pool of single cells) can help untangle their compositional and transcriptional makeup and differences – in particular, with regards to functional zonation, as well as clearance of apoptotic cells through an interplay of follicular subpopulations best resolved through imaging-based ST techniques.

### Keratins and immune-stromal microenvironment distinguish apical from basal and crypt from surface epithelium

Tonsil epithelium consist of two different types of actively differentiating epithelium: lining (surface) stratified squamous epithelium; and, reticulated crypts – deep invaginations of the tonsil in which the continuity of the outer surface epithelium is disrupted. The crypts represent a specialized compartment, important in the immunological functions of the tonsil, as they facilitate the interaction between immune cells and antigens, aiding in the recognition and removal of pathogens that enter the body through the oral cavity^[67]^. Both surface and crypt epithelial cells contain keratins, a protein family that forms intermediate filaments and maintains cell structure and function, contributing to mechanical strength and protection from stress. Keratin expression profiles depends on the type of epithelium, the differentiation status of the cells, and the biological context^[68]^.

To better characterize and map epithelial cells in the tonsils, we delineated 45 epithelial substructures comprising both crypt (all sections) and surface epithelium (adult sections only), based on the point pattern density of epithelial cells (C1 EPI01-06, A1 EPI01-10, and A2 EPI01-29; see Fig. 4a, Fig. S11, and Methods). These included about 152k epithelial cells (312k cells in total, also considering non-epithelial cells embedded in this layer). Across epithelial structures, we identified eight epithelial subpopulations that occupy distinct layers within the epithelium (from basal to apical axis) and were largely distinguishable by the keratin (KRT) transcriptional signatures (Fig. 4b-e): two apical (epi.api1/2), one basal (epi.bas) and suprabasal (epi.sup), as well as four transitional subpopulations (epi.trn1-4). These populations ordered from basal to apical represented differentiation states and were generally found across surface and reticular crypt epithelium (Fig. 4c,d). Basal epithelial (epi.bas) were characterized by NGFR and CD44 and represented the progenitor population^[69]^. At the protein level, PanCK increased from epi.trn1 to epi.trn4, and from epi.sup via epi.bas to epi.api1, but was erratic across sections for epi.api2 (Fig. 4f). To quantify the positioning of other subpopulations within the epithelium, we selected a structure that delineates the basal border of the epithelium (C1.EPI03), and estimated each cell’s distance to the apical boundary (Fig. 4g). Bm.cs and Bn IFN, ILC3 and Tc, as well as IgA+ and IgM+ PC were located near epi.bas, while Treg, The, Trm, NK/ILC1 subpopulations and activated naive B cells (Bn.act) were found more apically, followed by various myeloid subtypes and Bm; gran where distinctly apical.

**Figure 4:**
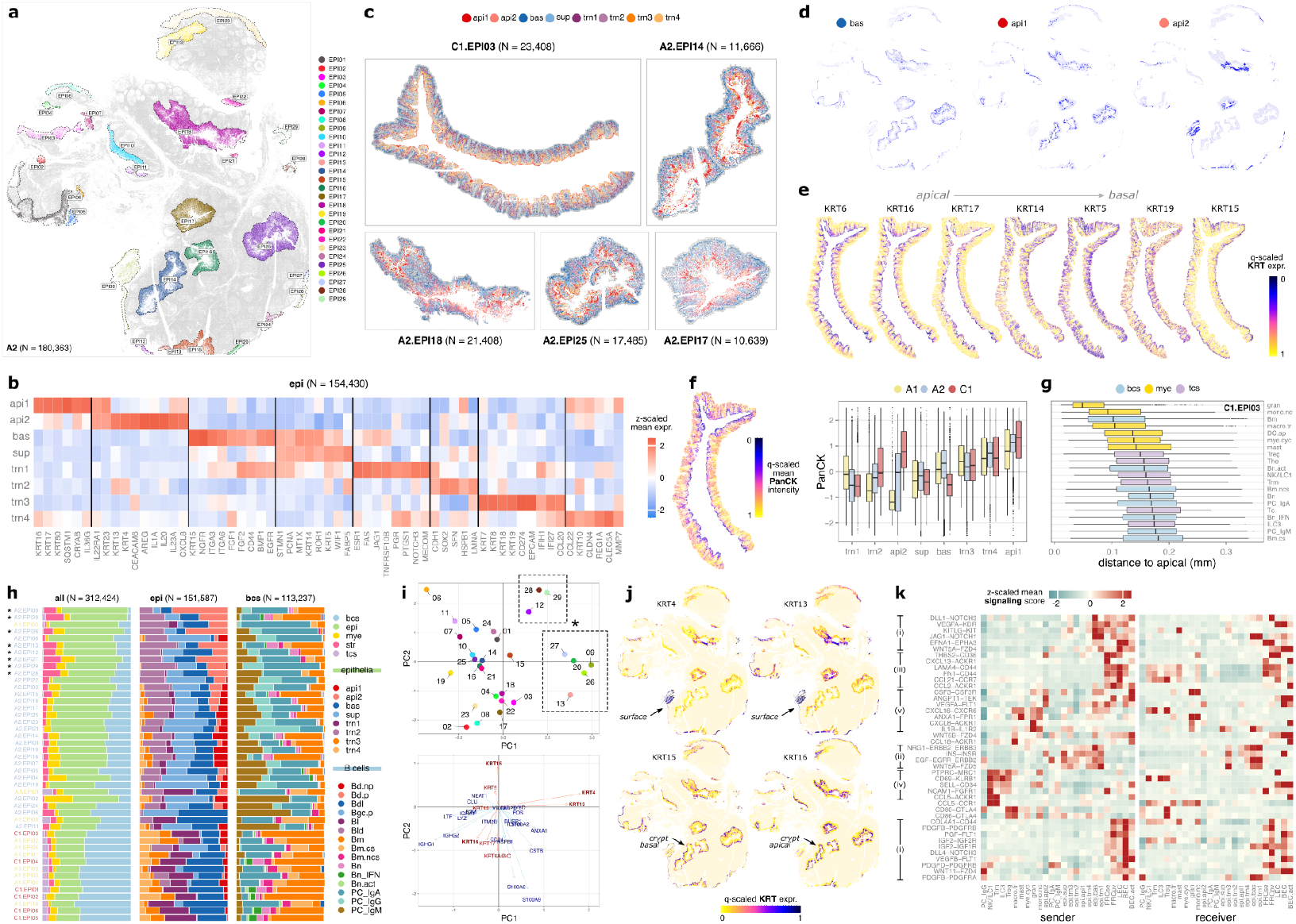
Epithelium. **(a)** Spatial plots highlighting programmatically identified epithelial substructures for section A2; dashed outlines are concave hulls of selected cells (cf., Fig. S11). **(b)** Heatmap of selected marker genes for epithelial subpopulations (data across all sections). **(c)** Spatial plots for selected substructures with cells colored by epithelial subpopulation, non-epithelial cells are grayed out. **(d)** Spatial plots highlighting basal (epi.bas) and apical (epi.api1/2) epithelial subpopulations; section A2. **(e)** Spatial plots with cells colored by the expression of selected keratin (KRT) genes; region C1.EPI03. **(f)** Spatial plot of mean PanCK intensities; region C1.EPI03. And, boxplot of intensities across epithelial subpopulations; x-axis ordered by median, y-axis values correspond to mean per-cell intensities, arcsinh-transformed with a cofactor of 200, and z-scaled by section. **(g)** Boxplot of cell-wise distances to the apical epithelial boundary, stratified by subpopulation; y-axis ordered by median; included are the 20 most frequent non-epithelial subpopulations. **(h)** Compartment and subpopulation frequencies for epithelial and B cell subpopulations across epithelial substructures; y-axis ordered by hierarchical clustering of left-most panel, labels colored by section. **(i)** First two principal components (PCs) and corresponding loadings (top-30 features) for pseudobulks of epithelial substructures (mean expression, including epithelial subpopulations only); section A2. **(j)** Spatial plots colored by selected keratinocytes (KRT) that exhibit surface (KRT4, KRT13), basal (KRT15), and apical (KRT16) expression; section A2. **(k)** Heatmap of cell-cell communication interactions between EPI/SM-related subpopulations (section C1); sender and receiver scores were averaged by subpopulation, ranked by fold change between sub-populations, and the top-50 selected for visualization; axes are ordered by hierarchical clustering of ‘sender’ values; left-hand side annotations indicate functionally distinct signaling modules.

Within EPI substructures, non-epi. subpopulations were mostly Bm and PCs, alongside some stromal and myeloid, and few T cells (Fig. 4h). Basal (bas) and suprabasal (sup) epithelial subpopulations were present across all substructures, while the two apical subpopulations (api1/2) tended to be mutually exclusive (cf., Fig. 4c,d). Transitional (trn) epithelial states were represented heterogeneously. To further investigate transcriptional differences between epithelial substructures – in particular, of surface and crypt types – we performed principal component (PC) analysis on their pseudobulk expression profiles (Fig. 4i). A group of substructures (12,28,29 and 09,13,20,26,27) was separated off by PC1, marked by high loadings on KRT4 and KRT13; these were opposed by immune-related genes (e.g., Ig’s). PC2 was driven by varying contribution of several KRTs and myeloid-related genes (e.g., S100A8/9), introducing variability between remaining substructures. The spatial distribution of KRT4 and KRT13 confirmed expression that is mostly localized to surface epithelium, while other keratins were expressed homogeneously in both crypts and surface types (Fig. 4j). Notably, these keratins were also specific to epi.api2 (Fig. 4b,d), which was more frequent in immune-depleted epithelial structures (Fig. 4h).

Across EPI structures and niches, CCC results captured diverse signaling modules favoring distinct functions (Fig. 4k): We identified abundant stromal homotypic and some heterotypic interactions (mainly with basal epithelial populations) aimed at sustaining the progenitor epithelial/stromal niche (e.g., NOTCH, WNT, IGF, PDGF, VEGF)^[70]^ – module (i). We also identified homotypic interactions to sustain epithelial proliferation and function (e.g., NRG, INS, WNT)^[71,44]^ – module (ii). Interestingly, PC IgA supported the survival and growth of epithelial cells through the EGF/EGFR axis.

Stromal populations were found to promote immune cell recruitment and adhesion through CCL21-CCR7, CXCL13-ACKR1 and LAMA4 FN1-CD44, among others – module (iii) – while Trms and ILCs engaged in communication with ECs and FRCs through CD69-KLRB1, SELL-CD34, NCAM1-FGFR1, CCL5-CCR1 ACKR1 for cell adhesion and retention – module (iv). Lastly, we identified a proinflammatory CCC module, largely mediated by cells of myeloid and mesenchymal origin on the apical side of the epithelium (e.g., CSF3, ANXA1, CXCL16, IL1B, CXCL8) – module (v).

Through a structure-based view, we dissected transcriptional and microenvironmental features of both apical/basal and, although less pronounced, surface/crypt epithelium, suggesting functional differences within coherent epithelium, as well as across epithelium at different tissue sites.

## Discussion

Single-cell sequencing has revolutionized biomedical research by enabling the transcriptome-wide analysis of phenotypic and functional heterogeneity at the level of individual cells. Recent spatial technologies have emerged from the convergence of single-cell transcriptomics approaches and advanced imaging techniques. These enable the detection of a large number of RNA molecules from tissues while preserving intact architecture^[72,73]^, thus allowing for the characterization of cellular interaction, organization and (dis)function in health and disease.

Yet, analyzing data from SMI remains challenging and lacks standardization compared to other single-cell omics assays, such as scRNA-seq^[7,8]^ and mass cytometry^[74,75]^. Library size-based normalization, for instance, has been shown to be problematic for targeted panels^[76,77]^. Many common quality control metrics (e.g., percentage of mitochondrial reads) do not apply. Failing to call spots causes false negatives (cf., dropouts in scRNA-seq), while false positives stem from probes misbinding or failure to strip. And, sample deteriorating over repetitive imaging-bleaching rounds can cause variability in detection efficacy depending on the preassigned barcodes and tissue-wide location of targets. Overall, the data is sparser and has a lower dynamic range (nearly binary); thus, it is unclear whether counts derived from imaging have the same statistical properties (e.g., mean-variance relationship) as sequencing-based count data.

Moreover, allocation of transcripts to cells relies on identifying cellular (membrane or nucleus) boundaries via segmentation. Various artifacts are associated with this process, such as spatial spillover, or ‘bleeding’, of transcripts^[78,79,80]^. Additionally, data are acquired through iterative imaging of predefined fields of view (FOVs). Lack of stitching during image pre-processing leads to border effects – fractured and/or duplicated cells being segmented near FOV borders with potential data inconsistencies. These effects were stronger in previous hardware and software versions of CosMx instruments, and have improved substantially with commercial updates. As of now, however, unnecessary cell loss and biases in spatial analyses that involve distance and neighborhood quantification remain.

Here, we addressed some of these issues by establishing a computational pipeline to analyze an ultimate of ~ 2M high-quality cells from 3 palatine tonsil sections – in a robust (e.g., quality control), scalable (i.e., parallelizable), versatile and reproducible fashion (using *Snakemake*^[81]^). Of note, standard scRNA-seq samples comprise 5-10k cells, which is on the order of cells that fall within few FOVs when applying imaging-based ST. While transcriptome-wide profiling of mammalian samples often yields data with dimensions on the order of 10^4^, the number of features required to (approximately) represent a single-cell omics dataset, its ‘intrinsic dimensionality’, is typically much lower. Thus, despite the limited number of target genes in our panel (~ 1000), we were able to identify 52 subpopulations across hematopoietic, epithelial and mesenchymal lineages, giving us an accurate representation of the cellular composition of the tonsil. The variability captured in target counts per area across different subpopulations reflected the adequacy of the panel to profile each of these, the target sensitivity for different cell types, and/or the differences in transcriptional complexity across subpopulations.

Moving to the microanatomical level, we describe seven niches or regions that compartmentalize the tonsils into functionally distinct units, in line with the literature^[9,10,3]^: the epithelium and submucosa provide a barrier from the outside world, while facilitating the recognition and removal of pathogens; follicles, comprised of germinal centers and their mantle, produce memory B cells and high-affinity antibody-producing PCs to fight recurrent exposures; the T cell zone supports trafficking of cells, T cell activation and differentiation; the B/T border represents an interface between compartments, where B and T cells interact; and, the connective tissue septum houses tissue-resident populations that monitor both local signals from tissue, and those entering from the lymphovascular system. Beyond what has been previously shown, we map subpopulations within and across these niches, and perform distance-aware modeling of inter-cellular signaling among them. Through this, we describe the tissue topology of several clinically relevant signaling pathways; e.g., TNF and GALECTIN have been linked to the pathobiology of different lymphomas^[28,29,30,64,65]^, and were pinpointed to the mantle zone in our study.

The enrichment described in BAFF and CD70 signaling at the T/B border is in line with their dual role promoting the development and activation of follicular and MZ B cells, as well as Tfh^[34,35]^. Cells from different lineages share not only tissue niches, but also expression patterns and signaling pathways, pointing to a high level of tissue compartmentalization, in terms of both cellular distribution and molecular fingerprints. This highlights the importance of molecular topology within tissues in relation to function (beyond cell topology in relation to function). Such molecular commonalities within tissue compartments are, however, overseen in experimental settings that lack spatial resolution, but might be crucial to gain a thorough understanding of how tissue microenvironments shape cell phenotypes and vice versa.

In addition, we delineated spatial units that allowed us quantify and compare the particular cell organization and gene expression patterns defining their architecture and function. On the one hand, comparative analysis of epithelial substructures let us delineate microenvironmental and transcriptional differences between apical/basal and crypt/surface epithelium. While the surface epithelium generally showed a consistent keratin (KRT) profile, KRT4 and KRT13 in particular, which are markers of mature squamous cell differentiation, the crypt epithelium exhibited a more variable KRT expression, in line with previous reports^[82]^. The reduction or absence of these KRTs in the crypts suggests a less differentiated epithelial state compared to the surface.

On the other hand, we demarcated individual GCs, letting us compare their composition and quantify expression gradients along a biologically well-defined physical direction: from light to dark zone (L/DZ), which is located near the GC mantle that is, in turn, dominated by naive B cells. Notably, our procedure differs conceptually from standard approaches to compare discrete LZ and DZ subpopulations (via differential gene expression analysis), or to identify spatially variables genes in an unsupervised fashion (via assessment of auto-correlation). Instead, we treat gene expression as a continuum, and leverage the LZ-DZ axis to pinpoint meaningful spatial trends.

The architecture of GCs is meant to allow for B cell proliferation, affinity maturation and, putatively, antibody diversification. Using class switch recombination (CSR) at the GC-level as a mark for maturation did not correlate with known cells and/or molecules driving this process. On the one hand, these results suggest that CSR occurs outside of follicles, complementing a previous murine study^[53]^. On the other hand, this could mean that Ig isotypes within GCs are inadequate as a temporal mark of follicles.

As they carry out their function, there are numerous cells undergoing apoptosis in GCs. Thus, DNA, together with organ specific self-peptides, could be presented to GCBCs through FDCs, which can lead to autoantibody production. In human autoimmune diseases, somatically hypermutated B cell receptors and autoantibodies are formed in GCs. We distinguish GC-resident macrophages that have been shown to engulf BC apoptotic bodies through dendrite-like protrusions^[58]^, with the crucial role of maintaining homeostasis in GCs. This resident population has been largely overseen and thus understudied in scRNA-seq analysis of SLOs. We could visually validate the identity of these macrophages based on their particular distribution in the tonsil, emphasizing the power of imaging-based ST to identify rare but relevant and spatially peculiar cell types.

TNFSF9 is expressed by GCBCs and has been postulated as a marker for memory B cells and lymphoma subtypes^[64,65]^. Its receptor, TNFSRF9 (4-1BB), is a classical T cell activation marker, which is also expressed by macrophages^[83]^ and FDCs^[22]^ costimulating B cells in GCs. This R/L pair engages in bi-directional communication^[84]^. In addition, loss of TNFSF9 in B cells has been linked to autoantibody production and the spontaneous formation of large GCs in mice^[66]^. We specifically detected GC B cells expressing TNFSF9 that signals through TNFSRF9 on FDCs and macro.tbm/act. Since we also observed these populations interacting directly within GCs, and based on the phagocytic function attributed to FDC-licenced TBMs, it is tempting to speculate that TNFSF9-TNFSRF9 signaling between the aforementioned follicular cell types represents an important homeostatic mechanism for GC maintenance, and a potential biomarker for follicle-associated diseases.

We argue that, unlike single-cell sequencing-based approaches, these data are a closer reflection of the true proportions of cell lineages within tissues, since there is no tissue digestion (physical or enzymatic) that can lead to loss of particular cell types; fibroblast, endothelial, and epithelial lineages, for instance, are not lost in favor of immune cells. In the context of SLOs in particular, FRCs and FDCs are key organizers of tissue architecture, and modulators of immune cell localization and function, so that their accurate representation is paramount. As such, sequencing-based studies of the past decade may have biased our understanding of tissue composition and complexity. SMI arguably has the potential to better resolve interactions that remain understudied at a transcriptional level, especially those where non-immune players are key (e.g., cancer-associated fibroblasts in the tumor microenvironment, aberrant epithelia in inflammatory diseases or colorectal cancer, etc.).

In closing, the work presented herein constitutes a comprehensive resource for the community i) by providing a spatial characterization of the human tonsil at various scales (molecular, cellular and niche); and, ii) by providing a blueprint for spatial-centric analyses that can be applied to diverse organs and biological questions – in the Spatial Immunology field and beyond.

## Data availability

CosMx SMI data have been deposited on Zenodo at the following DOIs. Flat files (FOV placement, count matrices, cell metadata, segmentation boundaries, single molecules) for both slides S1 (sections C1 and A1) and S2 (section A2) are located at 10.5281/zenodo.16782547. We also make available *SingleCellExperiment* (.rds) and *AnnData* objects (.h5ad; written using *zellkonverter* ^[85]^) for R and Python, respectively, as well as language-agnostic *alabaster* file artifacts (written using *alabaster*.*sce*^[86]^) that, for each section, gather analysis results; also included are reference profiles that were used for low- and high-resolution clustering. The underlying scRNA-seq data is available through the *HCATonsilData*^[4]^ package via Bioconductor’s *ExperimentHub*^[87]^. Napari inputs (Zarr stores of IF markers, FOV and segmentation labels, single molecules) are located at 10.5281/zen-odo.16782637 (slide S1), 10.5281/zenodo.16782795 and 10.5281/zenodo.16784436 (slide S2).

## Code availability

All analyses were run in R v4.5.1^[88]^ with Bioconductor v3.21^[89]^. The computational workflow was implemented using *Snakemake* v7.26.0^[81]^, with Python v3.11.3. Data were exported from AtoMx as flat files and handled using the *SingleCellExperiment* container^[7]^ with a delayed backend (using *HD5Array* ^[90]^). Cell segmentation polygon coordinates were written to .*parquet* and read into R for visualization using *arrow*. Data were visualized with *ggplot2* and *patchwork* using custom scripts, while IF-based plots were captured in *napari* with *viewer*.*screenshot(). Snakemake* workflow and underlying R code are accessible at https://github.com/HelenaLC/tonsilove, including session information on the software versions and operating system used throughout this study.

## Acknowledgements

We thank Bruker Spatial (f.k.a. NanoString) for providing section A2 in order to test the successful installation of a new CosMx instrument, and permitting us to use the correspondingly acquired data. H.L.C. acknowledges support by Swiss National Science Foundation (SNSF) grant number 222136. L.L.-C. is supported by the European Union’s Marie Skłodowska-Curie Actions (MSCA) Postdoctoral Fellowship program under grant agreement No. 101146541. H.H. receives funding from the European Union’s H2020 research and innovation program (848028), from the European Research Council (ERC) (810287), from the Ministerio de Ciencia e Innovación (MCI) (PID2020-115439GB-I00 and PLEC2021-007654), from the LaCaixa Foundation (HR22-0031 and HR22-0172), from the Generalitat de Catalunya through the Suport Grups de Recerca AGAUR (2021-SGR), from ERA-NET Neuron/Ministerio de Ciencia e Innovación (MCI) (PCI2022-133012), and from an ASPIRE Award from The Mark Foundation for Cancer Research and the Scientific Foundation of the Spanish Association Against Cancer. E.C. was supported by the “la Caixa” Foundation Health Research 2022 Program (CLLSYSTEMS LCF/PR/HR22/52420015 (HR22-00172); the Ministry of Science and Innovation PID2021-123054OB-I00, Generalitat de Catalunya Suport Grups de Recerca AGAUR (2021-SGR-01172). A.P.R. receives funding from the MCIN/AEI/10.13039/501100011033 and FSE+ (RYC2022-035848-I) and the MICIU/AEI/10.13039/501100011033/FEDER/UE (PID2023-148687OB-I00).

## Authors’ contributions

H.L.C. performed computational analyses, generated figures, and wrote computational methods. A.P.R. and H.L.C. interpreted the data and results, conceptualized and drafted the manuscript. S.G. aided in spatial feature reconstruction and corresponding downstream analyses. I.R. helped with histological and biological interpretation of tissue and data, respectively. L.L.-C., G.F., M.K., J.I.M.-S. and E.C. selected sections and aided in the interpretation of results. P.L. and M.R. performed CosMx SMI experiments; M.R. wrote experimental methods. H.H. acquired funding. All authors read and approved the final manuscript.

## Methods

### Clinical metadata and experimental processing

#### Sample collection and clinical details

All donors or legal guardians gave informed consent for their participation in this study, which was approved by the clinical research ethics committees of the Clínica Universidad de Navarra and of the Hospital Clínic de Barcelona (HCB/2018/0992). Samples were reviewed at the Hematopathology Unit of Hospital Clinic of Barcelona and showed reactive follicular hyperplasia with several degrees of germinal center expansions. No atypical cells in the epithelium, stroma or lymphoid compartments were seen in the cases (available clinical metadata for each section are summarized in Table S1). Tissues were fixed in formalin and embedded in paraffin according to standard pathology protocols.

#### Sample preparation and data acquisition

In situ hybridization was performed on five-micron tissue sections following the CosMx Manual Slide Preparation, MAN-10159-01, from Bruker. Section C1 and A1 were run in the CosMx SMI instrument (software version 1.1) as part of a 4x slide run, and section A2 as part of a 2x slide run. Slides were baked in an oven at 60 °C overnight. Tissue sections were dewaxed twice in xylene (Merck) for 5 min, twice in ethanol (Panreac Appliedchem) for 2 min and then baked at 60 °C for 5 min. Afterwards, they were subjected to a target retrieval step using CosMx Target Retrieval Solution (Bruker) by heating them at 100 °C in a pressure cooker for 15 min. After target retrieval, tissue sections were rinsed with DEPC-treated water (DEPC H2O, ThermoFisher), washed in ethanol for 3 min and dried at room temperature for 30 min. An adhesive incubation frame was placed around the tissue section. The tissue was then digested with CosMx Proteinase K at 3 ug/ml concentration for 15 min at 40 °C on the lymph node tissue and 30 min on the tonsil tissue. Tissue sections were rinsed twice with DEPC H2O, incubated in 1:100 diluted CosMx fiducials for 5 min at room temperature and washed with 1X PBS (ThermoFisher) for 5 min. After digestion and fiducial placement, tissue was fixed with 10% neutral-buffered formalin (EMS Diasum) for 1 min to maintain soft tissue morphology, washed twice with Tris-glycine buffer (0.1 M glycine (Sigma), 0.1 M Tris-base (FisherScientific), in DEPC H2O) for 5 min, and then washed with 1X PBS for 5 min. Fixed tissue was blocked with 100 mM Sulfo-NHS-Acetate (NHS-acetate, ThermoFisher) diluted in CosMx NHS-acetate buffer for 15 min at room temperature and washed with 2X saline sodium citrate (SSC, ThermoFisher Scientific) for 5 min.

CosMx™ Human Universal Cell Characterization Panel (RNA, 1000 Plex) probes (Bruker, PN: CMX-H-USCP-1KP-R) were incubated at 95 °C for 2 min and immediately transferred to ice. Probe mix was prepared according to CosMx protocol and then pipetted over tissue within a frame seal and an incubation frame cover on top. Hybridization took place at 37 °C for 18 hours. Following ISH probe hybridization, incubation frame cover was removed, tissue sections were washed twice in a buffer comprising 50% formamide (VWR) in 4X SSC at 37 °C for 25 min and twice with 2X SSC for 2 min each at room temperature. Next, slides were incubated with CosMx Nuclear Stain for 15 min at room temperature and washed in 1X PBS for 5 min. All tissue sections were incubated for 1 hour at room temperature with a mixture of CosMx™ Human Universal Cell Segmentation Kit (RNA) and CosMx™ Human IO PanCK/CD45 Supplemental Segmentation Kit (RNA). Samples HuN-29-0185 cut 03JUN2024 and HuN-29-0185 cut 03JUN2024 contained additionally CosMx Human CD68 A La Carte Marker, Ch5 (RNA), in the Cell Segmentation mix. After three washes with 1X PBS, the incubation frame was removed and a CosMx flow cell applied with the CosMx Flow Cell Assembly Tool. Slides were then ready to load on the CosMx SMI instrument according to CosMx Instrument User Manual MAN-10161-02 03.

The pre-bleaching profile for both runs was C; Cell Segmentation module D and A was selected for the 4x and 2x slide run, respectively. Both runs were set to 24s DNA, 100s Membrane and 33s PanCK exposure time. The 4x slide run had a 100s CD45 exposure time, while the 2x slide run was performed at 50s CD45 exposure time. The A La Carte Marker (CD68) on the 2x slide run was set at 25s exposure time. 200 fields of view (FOVs) were selected for each slide.

### Computational data processing and analysis

#### Quality control

Because lack of FOV stitching during cell segmentation can result in fractured and duplicated cells near FOV borders, we filtered out cells too close to any FOV border in order to mitigate potential artifacts in downstream analyses; Fig. S2a. Specifically, we used half the radius of an average (approximately circular) cell as a threshold on FOV border distance (according to 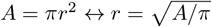, where *A* denotes the average cell area in a given section). For each section, we deemed as outliers cells below 3 median absolute deviations (MADs) in uniquely detecting RNA species, total RNA counts, and counts per area (computed on the log-scale using *scater* ‘s *isOutlier()* function^[91]^; Fig. S2b). Jointly, these criteria removed ~ 6% of cells per section, retaining 1,960,355 cells from 2,089,984 called segmented originally; Fig. S2c and Table S1. Henceforth, ‘expression’ values means log-transformed library size-normalized counts, computed with *scater* ‘s *logNormCounts()* function.

#### Semi-supervised clustering

Reference profiles were computed by summing counts per sample and cluster, library size normalization, and then averaging normalized counts across samples (using *scater* ‘s *aggregateAcrossCells()* for aggregation and *logNormCounts()* for normalization with *log = FALSE*). Semi-supervised clustering was performed using *InSituType*^[92]^ following the authors’ recommendations; briefly: To account for platform effects, reference profiles were adjusted using *updateReferenceProfiles()* with per-cell background estimated as the fraction of total RNA counts stemming from negative probes. Cohorts (via *fastCohorting()*) were based on mean immunofluorescence stains (PanMem, PanCK, DAPI, CD45 and, for section A2, CD68), as well as cell area and aspect ratio. Lastly, *insitutype()* was run with *n clusts* 6-12, yielding 15 *de novo* and 14 labeled clusters, which were annotated into 18 subpopulations.

Next, we stratified subpopulation into five compartments: B cell (Bn/m, Bd/l and PC; N=5), T cell (Th/c and ILC; N=3), myeloid (mono, DCc/p, macro, mono, mast and gran; N=6), stromal (EC, FDC and FRC; N=3) and epithelial (epi; N=1) subpopulations. These were subjected to another round of clustering using high-resolution reference profiles when available (myeloid, B and T cell compartment), and distinct feature selections for each subset. Specifically, we used *scran*’s *findMarkers* function (*direction=“up”*, blocking by donor) to test for differentially expressed genes within scRNA-seq clusters; features that ranked among the top-100 in any pairwise comparison, and which were present in our panel, were used for semi-supervised clustering. For reference-free compartments (epithelia, stromal), we selected for features that ranked among the top-400 in terms of fold change between average self-expression (i.e., EC/FDC/FRC for stromal; epi for epithelial) and all other subpopulations.

#### Niche analysis

To identify spatial context, we first quantified subpopulation frequencies among each cell’s local neighborhood; specifically, considering cells within a 20um radius (using *RANN*’s *nn2()* function with *searchtype=“radius”*). The resulting (cells × clusters) matrix of proportions, pooled across sections, was then used as input to *k*-means clustering with *k=10*. The resulting clusters were annotated into seven niches based on their subpopulation composition and spatial organization across tissue sections (Fig. 2a-c).

#### Cell-cell communication

Cell-cell communication analysis was performed using *COMMOT* ^[20]^, an optimal transport-based method that considers physical cell-to-cell distances and can account for competing sender/receiver signals as well as multimeter receptors. We considered 181 interactions (143 sender-receiver pairs from 38 pathways, 143 unique features) that were available through *CellChatDB*^[19]^, and for which all interaction partners were represented in our 980-plex panel. The average cell area in our data was ~ 48.71um^2^; here, we considered 20um as a threshold on signaling distance, which is about five times an average (approximately circular) cell’s radius 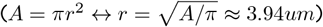.

#### Spatial feature reconstruction

We used the *sosta* R/Bioconductor package to identify epithelium (EPI) and germinal center (GC) structures following the authors’ recommended analytical steps: i) estimating reconstruction parameters (smoothing bandwidth, density threshold); and, ii) using these to identify individual structures. Specifically, we reconstructed structures based on the point pattern density of cells with a (low-resolution) subpopulation annotation of epi and Bd/l or FDC, respectively (argument *markSelect*).

Because epithelia were organized into extremly well-defined structures (i.e., present or absent), we multiplied the threshold estimate for EPI by 0.1 to better retain regions of comparatively lower density (e.g., at the tissue boundary). For GC structures, bandwidth estimates were multiplied by to obtain smoother structures; in addition, we estimated a concave hull around preliminary GCs and refined selection accordingly (using *concaveman* with *concavity=100*).

Filtered for structures comprising at least 200 epithelial and 100 Bd/l cells or FDCs, respectively, we obtained 6/10/29 EPIs (Fig. 4a, Fig. S11) and 45/26 GCs (Fig. 3a, Fig. S9) for sections C1, A1 (and A2). These were subjected to various downstream analyses (see next section).

#### Structure-level analyses

For each EPI structure, we computed pseudobulks (mean of library size-normalized and log-transformed counts) using cells with a low-resolution annotation of epi. These were subjected to principal component analysis (PCA) using the same set of features that had been used for subclustering of the epithelial compartment (Fig. 4i).

For each GC, we computed the median coordinates of Bn that fall within a 10um polygonal expansion from a structure’s (concave hull) boundary. Distances of within-GC cells to these reference points were quantile-scaled for each GC using lower and upper 1% quantiles as boundaries; i.e., distances are not comparable (in absolute terms) across GCs. As such, they reflect the relative positioning of cells within the GC, low/high distances being close/far from the mantle. To quantify corresponding gene expression gradients, we grouped cells into 27 bins of 0.04 distance, then averaged gene expression by bin and across GCs (Fig. 3d).

To score the maturity of GCs based on Ig isotypes, we performed a 20um negative expansion of their concave hulls as to mitigate bleeding near the mantle interface (of IgD and IgM in particular), and refine cell selections accordingly. Next, the expression of each Ig was ranked across GCs – by decreasing value for IgD and IgM, and increasing value for the remaining isotypes in the panel (IgG1, IgA1, and IgG2). Lastly, GC maturity was computed as the average rank across Ig’s. For example, a GC with (relatively) high IgD/M and low IgA/G expression will receive a low score -early-; vice versa, IgD/M-low and IgA/G-high GCs yield a high score -late-(Fig. 3g).

To investigate orthogonal readouts with respect to GC maturity, we quantified GC surface areas, their proportion of specific subpopulations (e.g., Tfh cells), as well as the average expression of relevant molecules (e.g., IL10) and genes associated with proliferation (e.g., MKI67) across Tfh and Bd cells, respectively. These, in turn, we visualize against GC maturity ranks (Fig. 3h).

