## Supplementary Material for "A transcriptional map of human tonsil architecture: beyond the sum of (single cell) parts"

| **Section** | **Slide** | **W x H** (mm) | **#FOVs** | **#Cells/FOV** | **#Cells** (raw\|fil.) | | **%** | **Sex** | **Age** | **Metadata** |
|---|---|---|---|---|---|---|---|---|---|---|
| C1 | 1 | 5.65 x 6.19 | 93 | 6,351 | 590,684 | 552,414 | 93.5 | M | 4 | Tonsillitis |
| A1 | 1 | 9.87 x 5.35 | 64 | 7,409 | 474,183 | 442,647 | 94.2 | M | 33 | Sleep apnea |
| A2 | 2 | 13.3 x 15.3 | 502 | 2,042 | 1,025,117 | 965,294 | 93.4 | n/a | n/a | n/a |
| | | sum/mean: | 659 | 3,171 | 2,089,984 | 1,960,355 | 93,8 | | | |

**Table S1: CosMx SMI data overview.** Sections were acquired in two runs with one slide each including two and one section(s), respectively. Width (W) and height (H) correspond to the largest difference in tissue-wide x- and y-coordinates, respectively. The scanning area amounts to $(512\text{um})^2$ per field of view (FOV), i.e., $659 \times (512\text{um})^2 \approx 1,73\text{cm}^2$ across sections. Cell counts correspond to the number of cells detected across all FOVs and those retained after filtering, respectively.

| **B & plasma cells** | |
|---|---|
| Bgc.p | proliferative germinal center B cell |
| Bd.p | proliferative dark zone B cell |
| Bd.np | non-proliferative Bd |
| Bdl | dark-light transitioning |
| Bl | GC light zone B cell |
| Bld | light-dark transitioning |
| Bn | naive B cell |
| Bn.act | activated Bn |
| Bn_IFN | interferon Bn |
| Bm | memory B cell |
| Bm.cs | class-switched Bm |
| Bm.ncs | non-class-switched Bm |
| PC_IgA/G/M | IgA/G/M+ plasma cell |

| **T cells** | |
|---|---|
| Tc | cytotoxic T cell |
| Tcn | cytotoxic naive |
| Th | helper T cell |
| Thn | naive Th |
| The | effector Th |
| Tfh | follicular helper |
| Trm | resident memory |
| Treg | regulatory |
| Tcyc | cycling |
| ILC3 | type 3 innate lymphoid cell |
| ILC1/NK | type 1 ILC / natural killer cell |

| **myeloid** | |
|---|---|
| gran | granulocyte |
| DCc | conventional dendritic cells |
| DCp | plasmacytoid DC |
| DC.ap | antigen-presenting DC |
| mye.cyc | cycling myeloid |
| mono.c | classical monocyte |
| mono.nn | non-classical mono. |
| macro.tr | tissue-resident macrophage |
| macro.act | activated macro. |
| macro.tmb | tingible body macro. |
| macro_ag.pres | antigen-presenting macro. |

| **stromal** | |
|---|---|
| BEC | blood endothelial cell |
| BEC.act | activated BEC |
| LEC | lymphatic EC |
| FDC | follicular dendritic cell |
| FRC | fibroblast reticular cell |
| FRCpv | perivascular FRC |
| FRCse | sub-epithelial FRC |
| FRCtcz | T cell zone FRC |
| FRCcts | connective tissue septum FRC |

| **epithelia** | |
|---|---|
| epi.bas | basal |
| epi.sup | suprabasal |
| epi.api1-2 | apical |
| epi.trn1-4 | transitional |

**Table S2: Glossary of subpopulation labels** including both low- and high-resolution annotations.

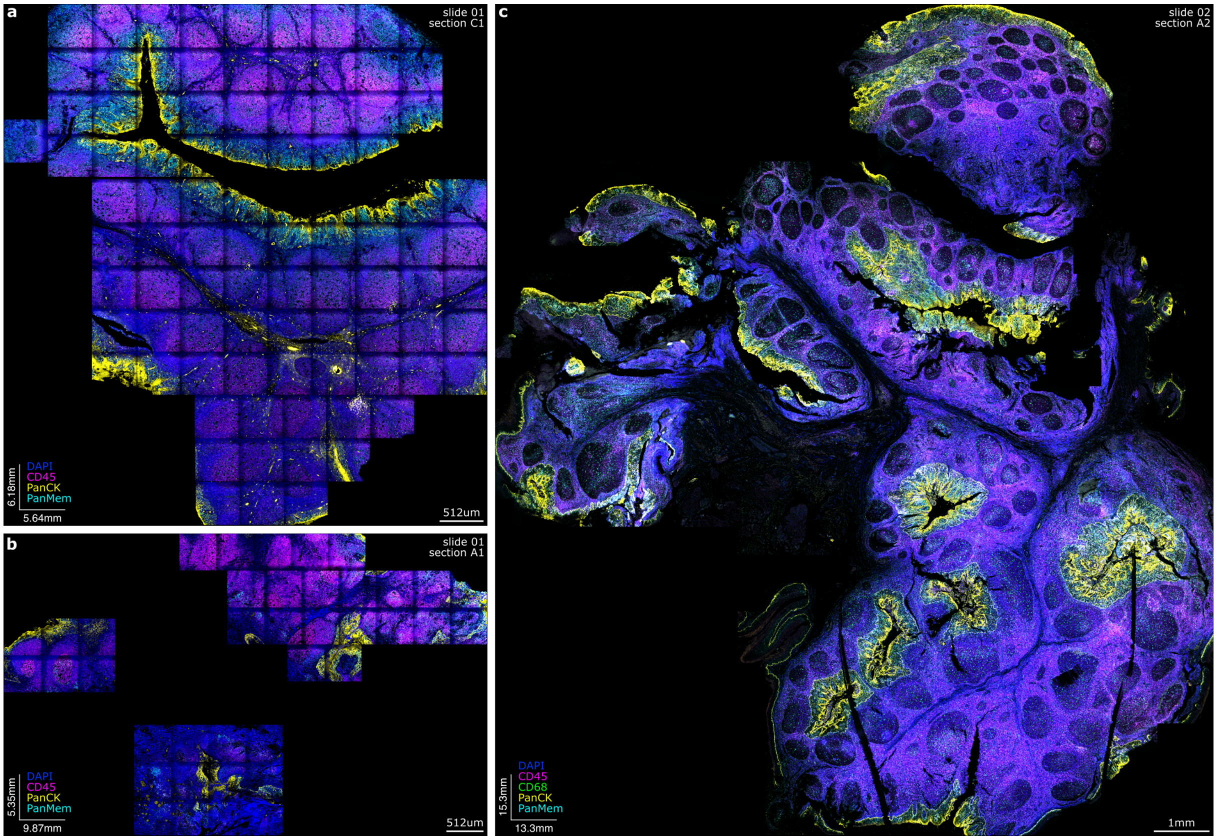


**Fig. S1: Immunofluorescence composite images** including DAPI (nuclei), PanCK (epithelia), PanMem(brane) and CD45 (immune). Data on sections C1/A1 **(a-b)** and A2 **(c)** were acquired on separate slides, the latter also including CD68 (macrophages). Scales bars in (a-b) correspond to the field of view (FOV) width/height. Overall section dimensions are also included; c.f. Table S1.

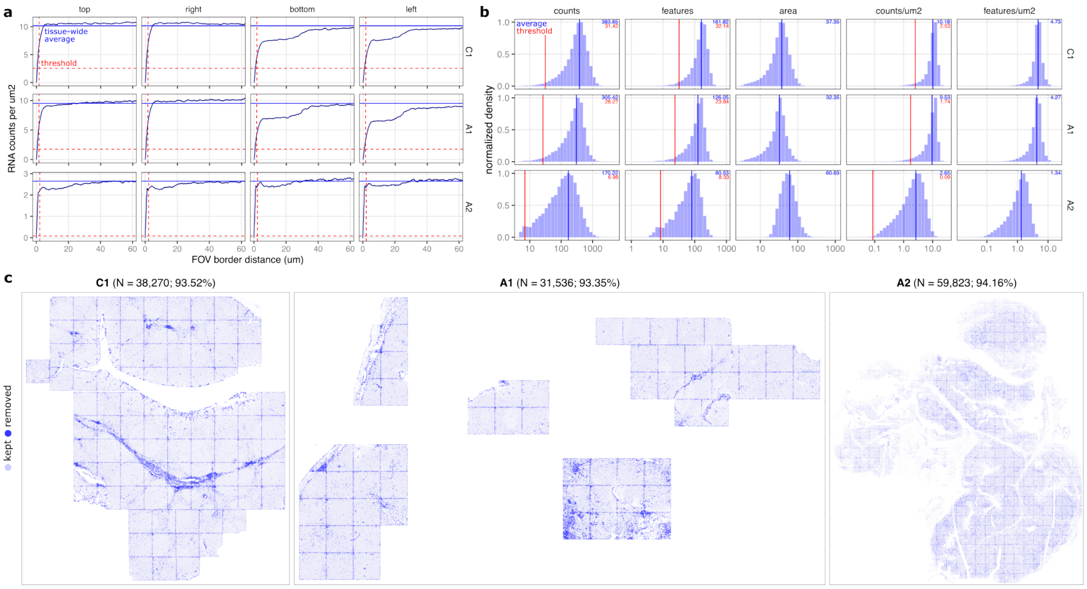


**Fig. S2: Quality control.** **(a)** RNA transcript counts per area (um$^2$) vs. distance (um) to field of view (FOV) borders; x- and y-axis values correspond to rolling means (window size 20); red lines indicate respective lower-bound filtering thresholds, blue lines indicate tissue-wide averages. **(b)** Histograms of (fltr) total RNA counts, uniquely detected features, and cell area (um$^2$), as well as total counts and unique features normalized for area; non-normalized x-axis values are $\log_{10}$-transformed; red and blue vertical lines indicate lower-bound filtering thresholds and medians, respectively. **(c)** Spatial plots highlighting low-quality cells.

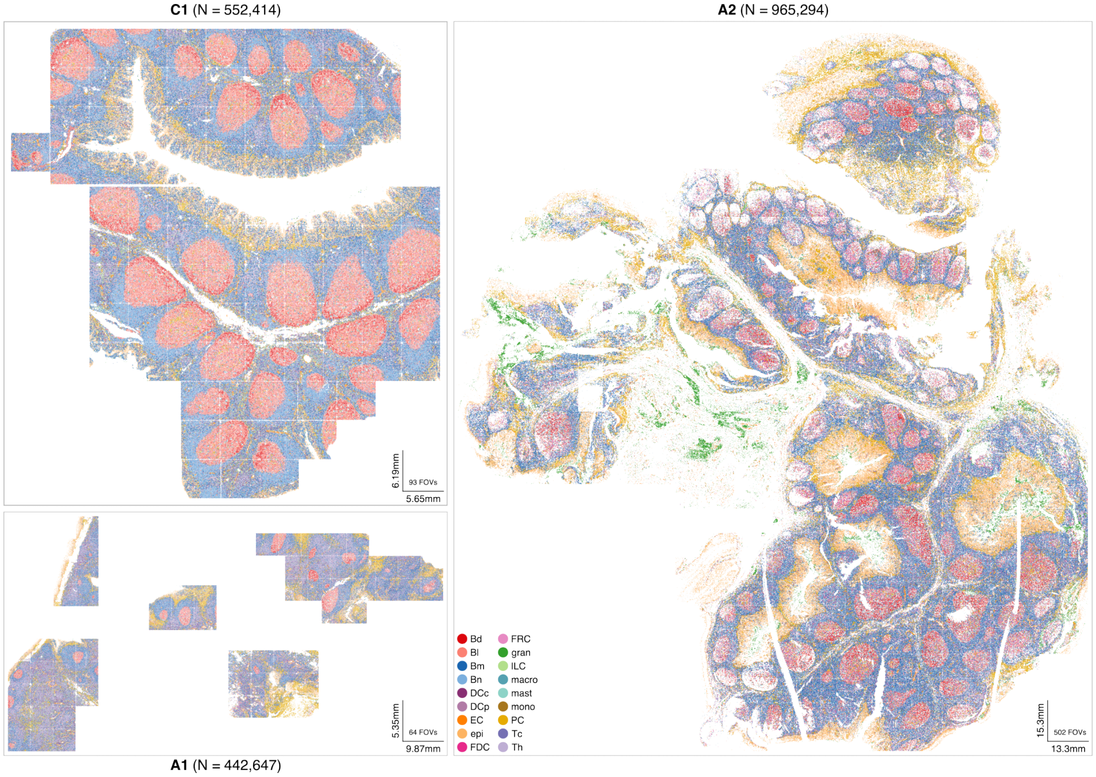


**Fig. S3:** Spatial plots with cells colored by low-resolution subpopulation assignment, including number of fields of view (FOVs) as well as physical dimensions (tissue-wide range of x- and y-coordinates) for each section.

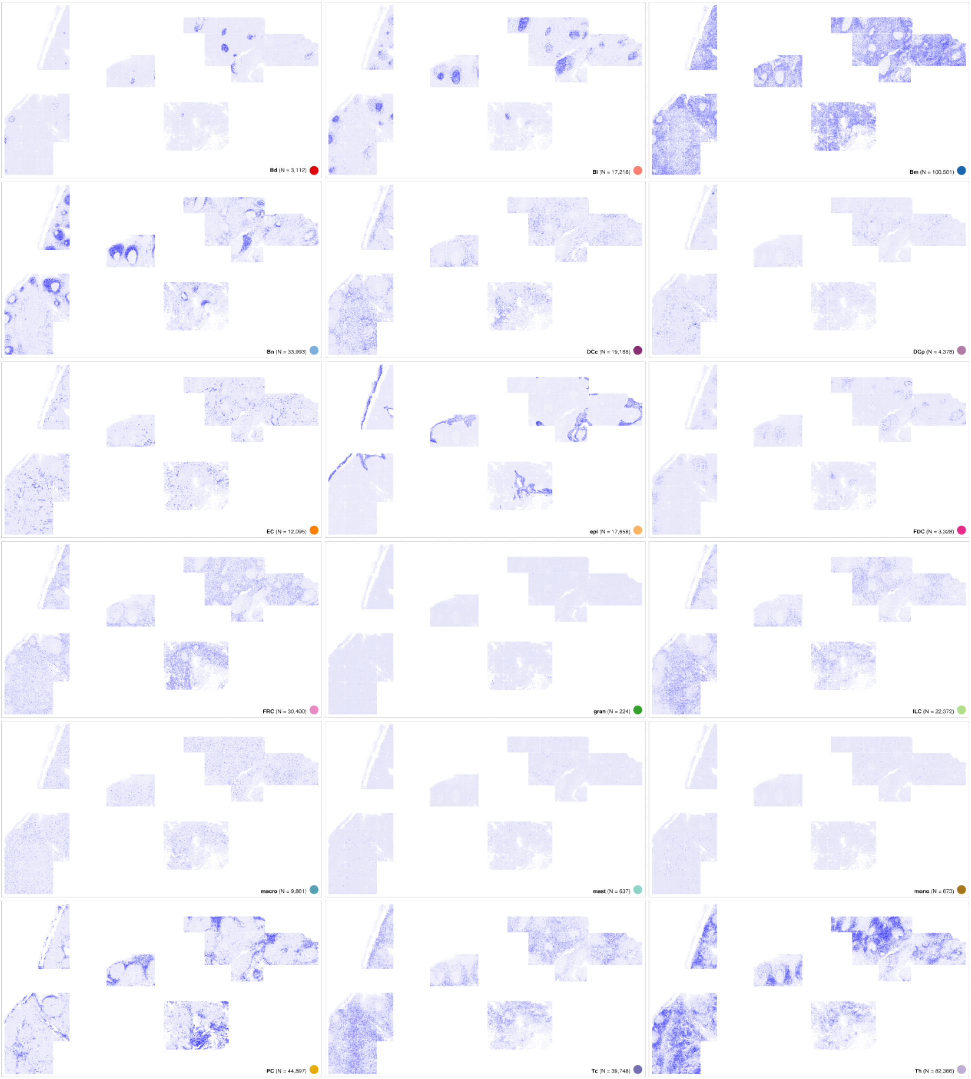

**Fig. S4:** Spatial plots highlighting individual low-resolution subpopulations; section A1.

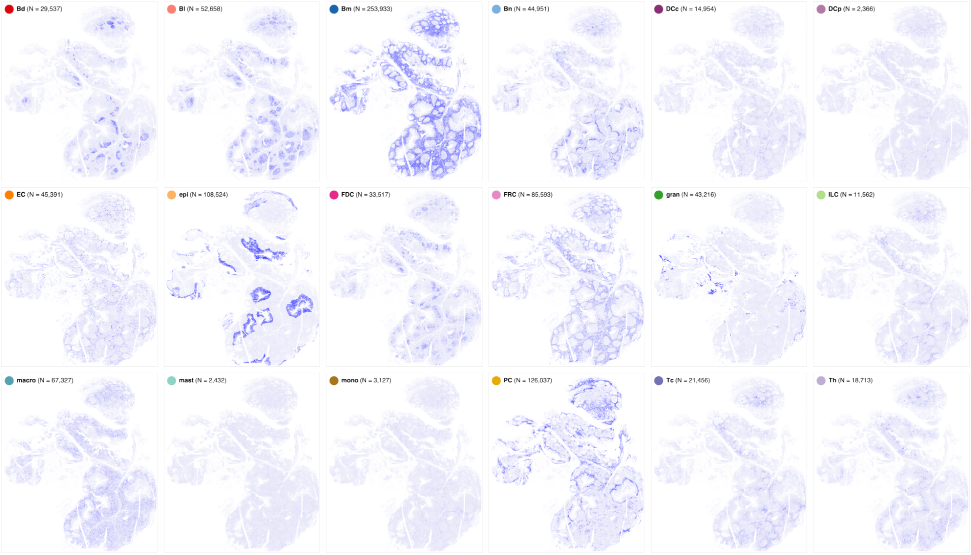


**Fig. S5:** Spatial plots highlighting individual low-resolution subpopulations; section A2.

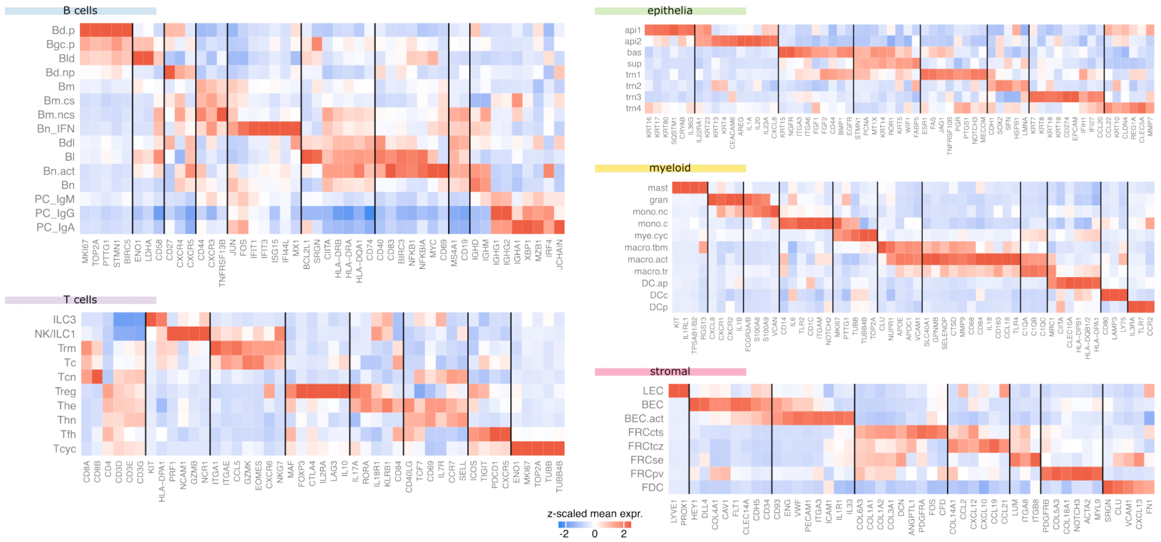


**Fig. S6:** Heatmaps of selected marker genes for each compartment (data across all sections).

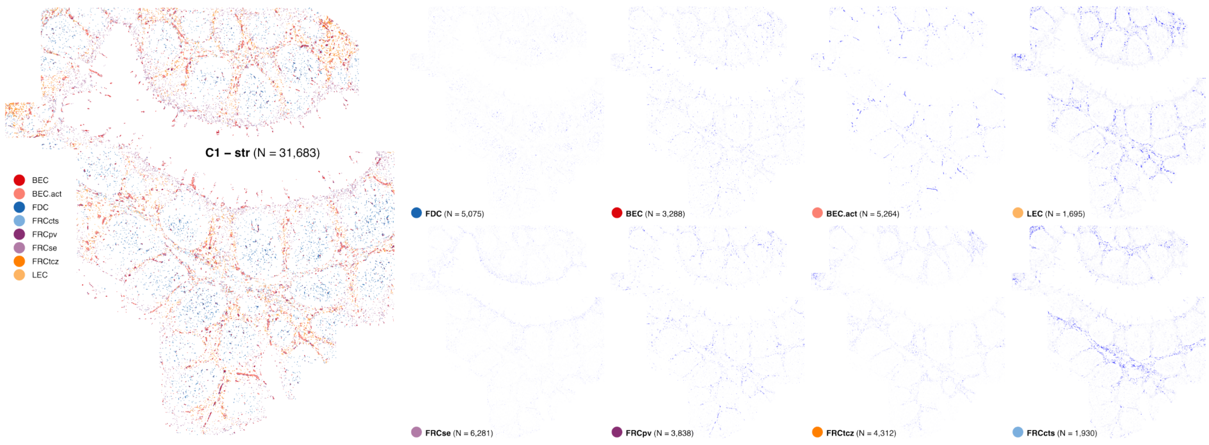


**Fig. S7:** Spatial plot colored by and highlighting individual stromal subpopulations, respectively; section C1.

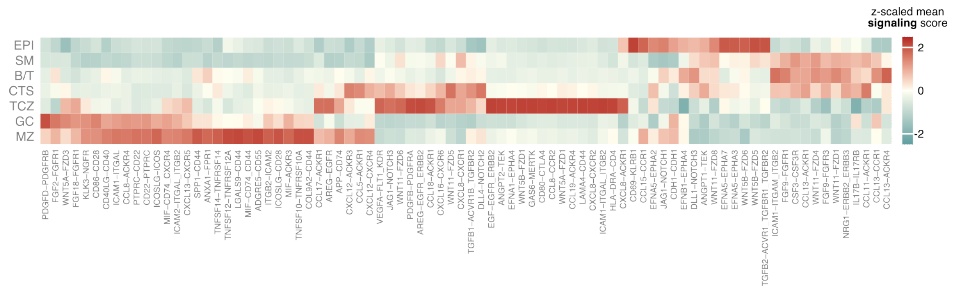


**Fig. S8:** Heatmap of cell-cell communication interactions across niches; values correspond to sender and receiver signals averaged across all cells in a given niche, averaged with each other, and z-scaled across niches; shown are the top-15 ranked interactions (in terms of fold change between niches).

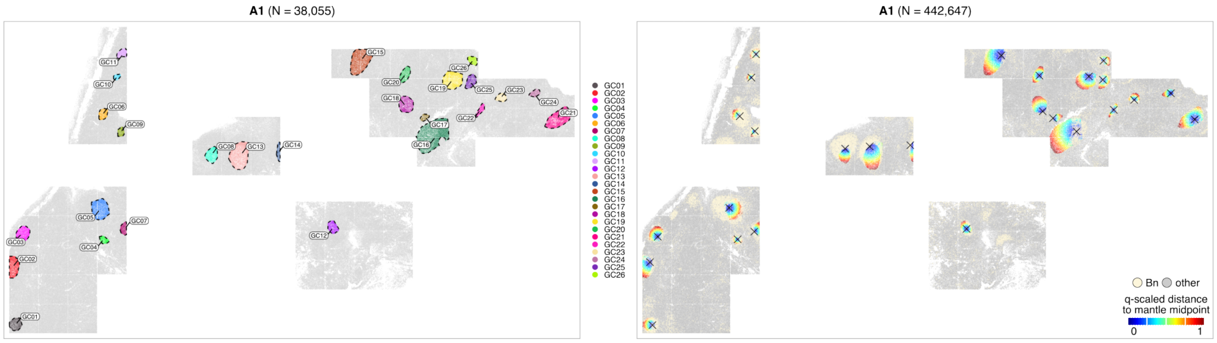


**Fig. S9: Germinal centers; section A1.** **(a)** Spatial plot highlighting programmatically identified germinal centers (GCs); dashed outlines are concave hulls of selected cells. **(b)** Spatial plot with cells colored by their relative distance to the median coordinates (black crosses) of Bn (off white) that fall within a 50um polygonal expansion from a given GC; cells that are neither Bn nor in any GC are grayed out.

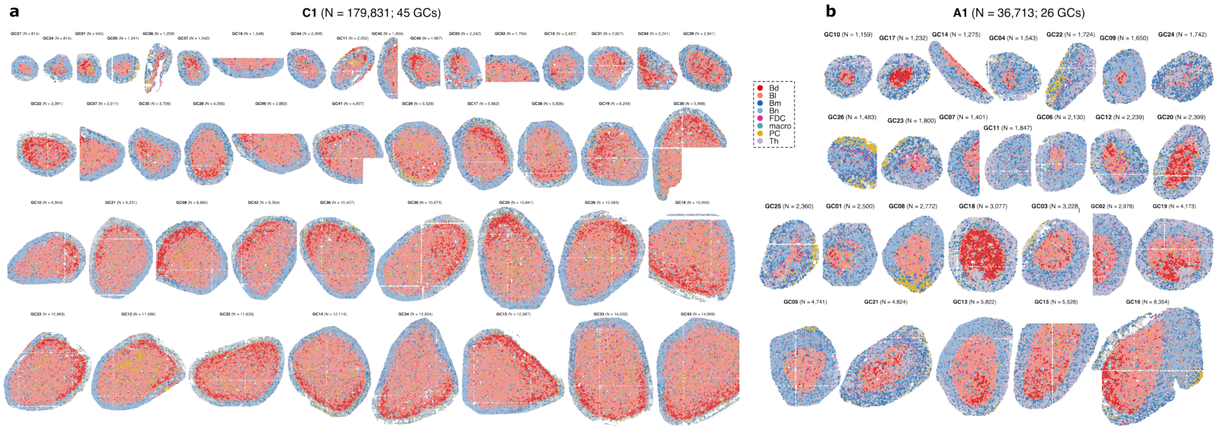


**Fig. S10:** Spatial plots of programmatically identified germinal centers (GCs) for sections **(a)** C1 and **(b)** A1; included are cells, colored by key subpopulations, that fall within a 50um polygonal expansion of the concave hull around selected cells; GCs are ordered by number of cells without expansion.

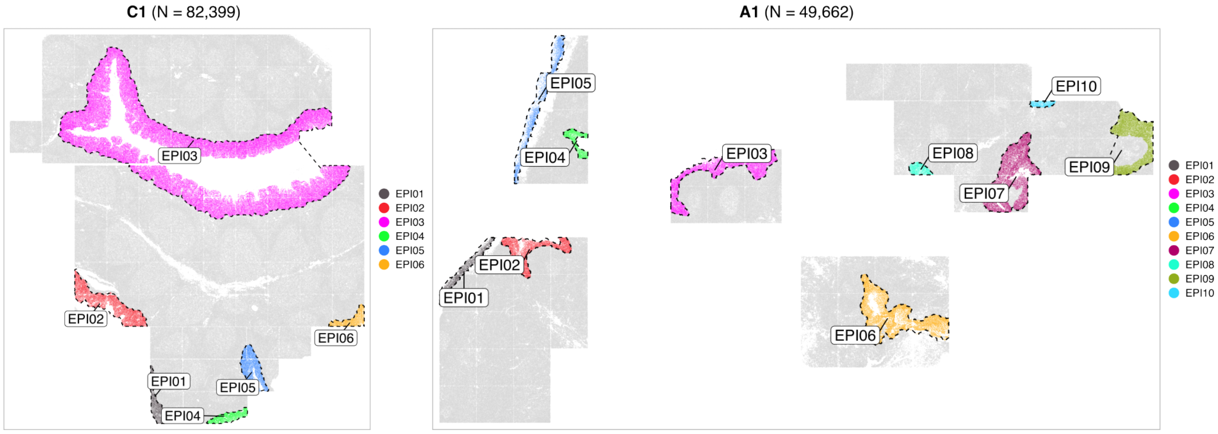


**Fig. S11: Epithelium; sections C1 and A1.** Spatial plot highlighting programmatically identified epithelial substructures (EPI); dashed outlines are concave hulls of selected cells.
